# NO VEIN is an ancestral immunity protein involved in plant-virus interaction

**DOI:** 10.64898/2026.09.30.755639

**Authors:** Eirik Lågeide, Maria Abraham, Nicolas Glück, Helena Pfingst, Lukas A. Burkhardt, Laura Medina-Puche, Rosa Lozano-Durán, Gal Ofir

## Abstract

The conservation of immune systems from prokaryotes to eukaryotes can guide discovery of immunity genes across the tree of life. Most studies to date have focused on metazoan systems, while the extent of ancestral immunity conservation in plants remains underexplored. Here we show that NO VEIN (NOV), a plant protein previously characterized for its essential developmental roles in *Arabidopsis thaliana*, shares domain composition and structural similarity with Tiamat, a bacterial antiphage protein. In light of the antiviral function of the bacterial protein, we asked whether NOV is involved in plant–virus interactions. We found that *NOV* knockdown in the model plant *Nicotiana benthamiana* resulted in an impaired transcriptional response to tobacco rattle virus, including reduced induction of defense genes, and observed increased potato virus x accumulation in a secondary infection. We also reveal that NOV, previously considered plant-specific, has homologs in animals - pointing to a shared origin. Together, these findings uncover a deeply conserved protein involved in plant-virus interaction, and demonstrate the power of prokaryote–eukaryote homology to reveal new aspects of plant biology.

## Introduction

All living cells, both prokaryotes and eukaryotes, must defend themselves against parasites and pathogens. Recently, multiple eukaryotic proteins involved in immunity were discovered to be homologous to prokaryotic immune systems (Aravind et al., 2024; Bernheim et al., 2024; Ledvina & Whiteley, 2024). Several notable examples include the conservation of Toll/interleukin-1 receptor (TIR) domain immune signalling (Essuman et al., 2022; Ofir et al., 2021), the widespread use of STAND-NTPase containing proteins in immunological processes (Chou et al., 2023; Contreras et al., 2023; Gao et al., 2022; Kibby et al., 2023), Argonaute-based RNA interference (Bobadilla Ugarte et al., 2023), cyclic-nucleotide immune signalling by cGAS-like enzymes (Hobbs & Kranzusch, 2024), and the conserved antiviral activities of Viperin enzymes (Shomar et al., 2024). Once explored, such homology can guide the discovery of new immunity-related genes. This approach was recently demonstrated by the discovery of new antiviral activities of human genes based on their homology to bacterial immune systems (Bonhomme et al., 2025; Cury et al., 2024), and by the discovery of new bacterial antiviral genes based on their homology to eukaryotic immune systems (van den Berg et al., 2024).

In many cases of ancestral immune gene conservation, the highly conserved portions of the homologous defense modules are typically limited to catalytic domains embedded within a protein whose auxiliary domains rapidly diversify and demonstrate domain modularity (Bernheim et al., 2024; Culbertson & Levin, 2023). To date, most such ancestral immunity modules were described in metazoans (Bernheim et al., 2024; Mordret et al., 2026).

Plant immunity involves several ancestral immune modules such as TIR domains, Argonautes and STAND-NTPase NLRs. However, multiple ancestral immune systems conserved in metazoans were not identified in plants to date, e.g. cGAS-like and Viperin enzymes (Bernheim et al., 2024). As knowledge of prokaryotic immunity is rapidly expanding, new opportunities arise to detect novel homologies between plant genes and recently-discovered prokaryotic immune genes. Unified search tools such as DefenseFinder and DefensePredictor can facilitate such searches (DeWeirdt et al., 2026; Tesson et al., 2022).

In this study, we identified the NO VEIN (NOV) protein of the model plant *Arabidopsis thaliana* to share domain composition and structure with the multi-domain bacterial immune protein Tiamat (Millman et al., 2022). Inspired by the antiviral function of Tiamat, we tested the effects of post-transcriptional silencing of the *Nicotiana benthamiana* homolog of *NOV* and observed an increased viral accumulation of potato virus X (PVX), indicating involvement in plant-virus interaction. Transcriptome sequencing of *NOV* knockdown plants demonstrated dampened induction of defense genes in response to tobacco rattle virus, suggesting an increased susceptibility to secondary infections. Beyond plants, we identify NOV homologs across multiple animal lineages, pointing towards a shared origin of plant and animal NOV proteins. Our findings identify NOV as a deeply conserved protein involved in plant response to viral infection, and demonstrate that deep conservation can serve as a guide to expand our knowledge of plant immunity.

## Results

### NO VEIN shares structural similarity and domain composition with the bacterial antiphage protein Tiamat

To search for plant homologs of bacterial defense systems, we applied the DefenseFinder tool on the *Arabidopsis thaliana* Araport11 proteome (Supplementary Table 1A-B) (Reiser et al., 2026; Tesson et al., 2022). This search identified the expected strong alignments between the prokaryotic pAgo and the plant Ago proteins (Bobadilla Ugarte et al., 2023). Most hits aligned to bacterial defense proteins over short, single-domain regions corresponding to common catalytic folds. Strikingly, the 2729 amino acid long protein product of the *A. thaliana NO VEIN* (*NOV*) gene (AT4G13750), a previously characterized plant developmental factor (Tsugeki et al., 2009), aligned to the ∼1500 amino acid single-protein bacterial defense system Tiamat (Millman et al., 2022) across most of Tiamat’s HMM profile (profile coverage 98.5%, e-value 7.2×10^−92^, matched across amino acids 1159-2709 in AtNOV) (Figure 1A, Supplementary Figure 1A-B). The mechanism of action of Tiamat in bacteria is unknown. Tiamat homologs were previously detected in Eukaryotes and Promethearchaeaceae in a general search for homologs of bacterial immune systems (Leão et al., 2024).

**Figure 1:**
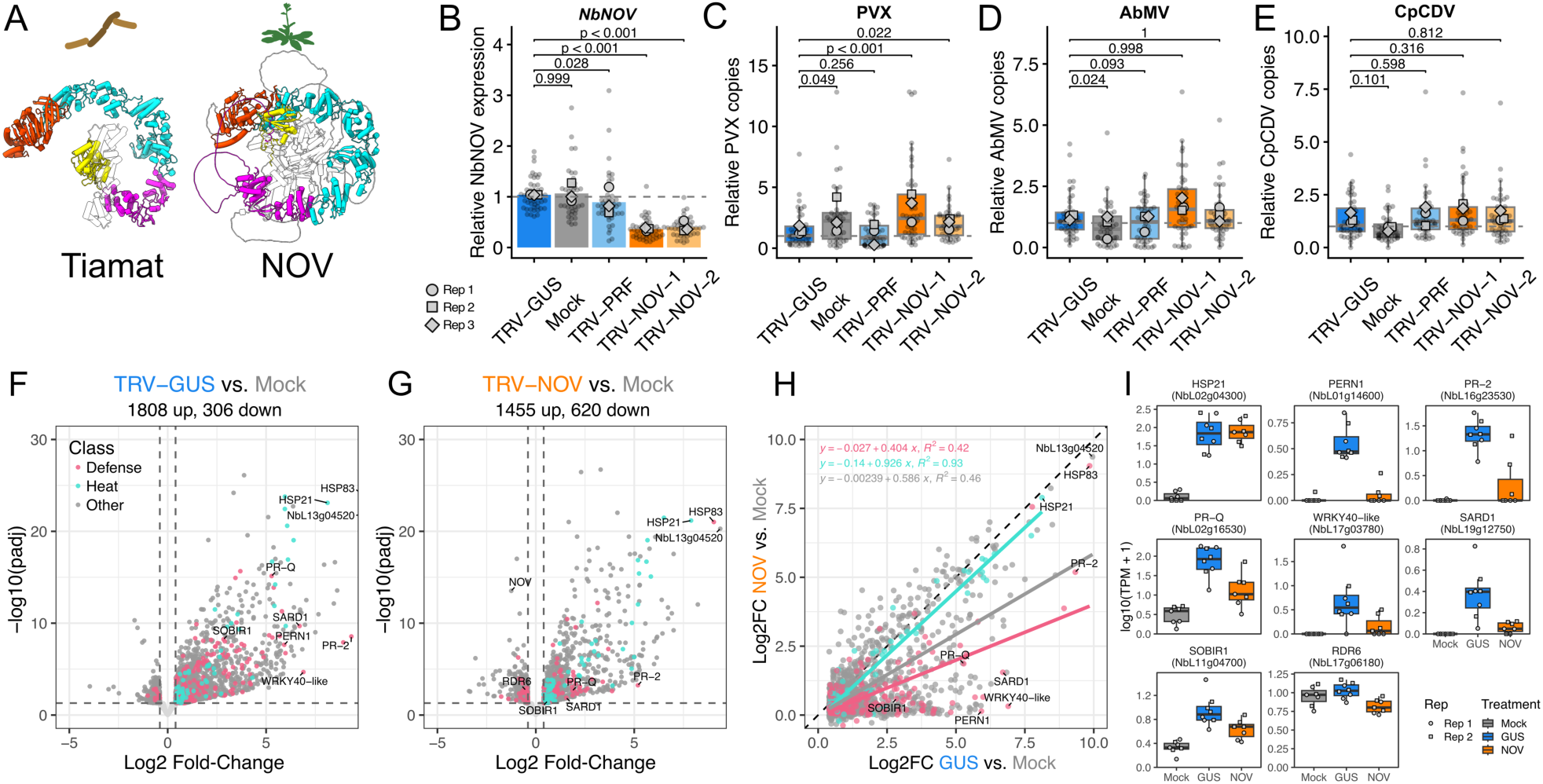
*NbNOV* knockdown affects viral infection and immune gene induction in *N. benthamiana*. **A** NOV and Tiamat share extensive structural similarity and domain composition - ATPase (orange), solenoid repeats (cyan), a beta-sheet motif surrounded by helical repeats (magenta) and NOV_C nuclease domain (yellow). AlphaFold3 predicted structures of *Bacillus cereus FORC_005* TmtA Refseq WP_046392403 and *A. thaliana* NOV AT4G13750 are shown. **B** *NbNOV* relative expression levels in the shoot apex of TRV- and PVX-coinfected plants 20 days post TRV inoculation, quantified using RT-qPCR, normalized to the NbeIF4A expression level in TRV-GUS infected plants (See Methods, and Supplementary Figure 2D-G for raw Ct values). Mock - no TRV, TRV-GUS - no-target negative control, TRV-PRF - targeting negative control, TRV-NOV-1 & TRV-NOV-2 - non-overlapping constructs targeting *NbNOV*. Results from 3 independent experiments, each including 12-16 plants per treatment, each datapoint represents an individual plant. The means of each independent experiment are marked with different shapes. **C-E** Normalized viral accumulation in *NOV* knockdown plants and controls. Plants were inoculated with TRV-VIGS constructs, and subsequently infected with PVX **(C)**, AbMV **(D)** or CpCDV **(E)** 10 days post-TRV inoculation. Viral accumulation was measured by qPCR in shoot apex following additional 10 days - a total of 20 days after the initial TRV inoculation (see Methods). **F-G** Volcano plot of differentially expressed genes in TRV-GUS infected plants vs. mock infection **(F)** or TRV-NOV infected plants vs. mock infection **(G)**. Red - Defense related genes, Cyan - Heat shock related genes **H** Correlation of fold-change upregulation level in TRV-GUS vs TRV-NOV. Each dot represents a single gene, significantly upregulated in response to TRV-GUS infection at p < 0.05 and absolute Log2 Fold Change > 0.4 **I** Expression levels (Transcripts per million) of selected differentially expressed genes highlighted in **(F-G)** across treatments.

We predicted the structures of AtNOV (pTM 0.42) and Tiamat (pTM 0.5) using AlphaFold 3 (Abramson et al., 2024) and found that the region corresponding to amino acids 1159–2709 of AtNOV shows extensive structural similarity to the entirety of Tiamat (Figure 1A, Supplementary Figure 1A). The shared region contains three conserved domains: a GHKL-type SACS ATPase (PF25794) at the N-terminus, a long alpha-solenoid repeat region with high structural similarity to sacsin internal repeats (Anderson et al., 2010), and a NOV_C PD(D/E)XK nuclease at the C-terminus (PF13020). The ATPase active site residues required for GHKL-type ATP hydrolysis and phage defense are conserved between NOV and Tiamat (Supplementary Figure 1B-C) (Millman et al., 2022). The C-terminal nuclease of NOV retains the catalytic residues characteristic of canonical PD(D/E)XK nucleases (Supplementary Figure 1B,D) (Steczkiewicz et al., 2012). We used EMERALD-UI (Grigorjew et al., 2023; Preoteasa et al., 2026) to quantify the robustness of the AtNOV^1156-2729^ – Tiamat structural alignment across an ensemble of near-optimal alignments (Supplementary Figure 1E). Of the aligned positions, 44% (694/1574) were classified as alignment-safe, i.e. robustly aligned across essentially all near-optimal alignments, distributed among 87 safety windows that are concentrated within the three shared domains.

### *NOV* knockdown increases potato virus X accumulation in *Nicotiana benthamiana*

Since Tiamat functions in antiphage defense in prokaryotes, we asked whether NOV has a role in host-virus interaction in plants. We identified a single *NOV* homolog in the genome of *Nicotiana benthamiana* (NbL01g24040, 92% coverage, 46% identity to AtNOV, Supplementary Figure 1C-D), a common model for plant-virus interaction studies. We knocked down *NOV* in *N. benthamiana* using tobacco rattle virus-induced gene silencing (TRV-VIGS) (Liu et al., 2002). In these experiments, a piece of the target gene is cloned into the TRV genome, leading to a collateral systemic RNAi-based post-transcriptional silencing of the gene of interest due to plant recognition of double stranded RNA formed during TRV replication (Shi et al., 2021). We used two independent knockdown constructs targeting non-overlapping regions of *NbNOV* (TRV-NOV-1 and TRV-NOV-2, see Methods). As a no-target negative control, a TRV-GUS construct targeting a portion of the *E. coli* β-Glucuronidase gene (GUS) was used. As an endogenous-target negative control, an NLR gene (NbPRF - NbL08g09580) (Dong et al., 2025; Kim et al., 2002), not known to be involved in viral immunity, was used. As an uninfected mock control, plants were infiltrated with *Agrobacterium* not carrying a viral construct. RT-qPCR performed 10 days following TRV infection confirmed significant reduction of *NbNOV* transcript levels to approximately 40% of the TRV-GUS control for both TRV-NOV-1 (p < 10^−4^) and TRV-NOV-2 (p < 10^−4^) (Supplementary Figure 2A). *NOV* knockdown had no major phenotypic consequences on plant growth under the experimental conditions used beyond the moderately stunted growth and mild hypersensitive response in lower leaves observed in all TRV infected plants (Supplementary Figure 2B).

Ten days after TRV inoculation, we re-infected plants with one of three viruses: the positive-sense ssRNA virus potato virus X (PVX), and the ssDNA geminiviruses chickpea chlorotic dwarf virus (CpCDV) or abutilon mosaic virus (AbMV). Viral accumulation at the shoot apex was quantified by RT-qPCR for PVX or qPCR for geminiviruses ten days after inoculation. We verified that *NOV* remained knocked down at the experimental endpoint (Figure 1B). *NOV* knockdown led to a significant increase in PVX copies for both *NOV*-knockdown constructs compared to the TRV-GUS negative control (NOV-1: 2.25-fold, 95% CI [1.42, 3.58], p < 0.001; NOV-2: 1.7-fold, 95% CI [1.06, 2.72], p = 0.022) (Figure 1C), while TRV copies were not significantly affected (Supplementary Figure 2C). The change in AbMV and CpCDV accumulation between *NOV-*silenced and control plants was not statistically significant (Figure 1D-E), indicating different effects of *NOV* downregulation on the interaction with different plant viruses.

### *NOV* knockdown impairs defense gene induction

To assess if the effect of *NOV* knockdown on viral infection was likely to be direct or indirect, we compared the transcriptome of *NOV* knockdown (TRV-NOV-1), control (TRV-GUS) and mock-infection plants 10 days post inoculation (see Methods). Infecting plants with the control TRV-GUS resulted in major transcriptional changes typical for virus infection, such as strong induction of heat shock, defense-associated, pathogenesis-related (PR), and systemic acquired resistance genes (Figure 1F, Supplementary Table 2) (Sahi et al., 2016; Whitham et al., 2006). We verified using the transcriptomic data that *NOV* expression itself was not affected by TRV-GUS infection (Supplementary Figure 3A).

The magnitude of the abovementioned transcriptional response was noticeably lower in TRV-NOV infected plants. This included fewer significantly upregulated genes (1455 vs 1808), more significantly downregulated genes (620 vs 306) and lower overall induction levels of upregulated genes (Figure 1G, Figure 1H). When comparing TRV-GUS and TRV-NOV infections directly, only 319 genes are significantly differentially expressed (94 upregulated, 225 downregulated, Supplementary Figure 3B). Among the genes detected as significantly downregulated in this comparison, 65% (148/225) were genes that were upregulated in TRV-GUS vs. mock infection, indicating a strong effect of *NOV* knockdown on the induction of viral infection response genes. Of these viral response genes, the reduced induction was more pronounced for genes with Gene Ontology (GO) “Defense response” annotation (Figure 1H). This was not the case for induced genes with GO “Heat response” annotation - whose induction level remained similar in TRV-GUS and TRV-NOV infected plants (Figure 1H). As viral expression levels were similar between TRV-GUS and TRV-NOV infected plants, these transcriptional differences do not seem to be the results of different TRV loads (Supplementary Figure 3C). Principal component analysis of the total transcriptome did not detect large-scale differences in the transcriptomic state of the plants between TRV-GUS and TRV-NOV treatments, indicating that the detected differences are specific to the effect of *NOV* knockdown (Supplementary Figure 3D). This suggests a specific involvement of NOV in the induction of defense-related genes of *N. benthamiana* in response to viral infection by TRV.

To better understand the effect of *NOV* knockdown on the expression of known immunity-related genes during TRV infection, we compared the expression of a strongly induced heat-shock related gene *HSP21* (O82013, AT5G12020) and six attenuated immunity genes, identified by their homology to Uniprot and TAIR entries: a PR2-family β-1,3-endoglucanase (P23547, AT4G16260), the PR3-family chitinase *PR-Q*, a *WRKY40*-like transcription factor (Q9SAH7, AT1G80840), the salicylic acid biosynthesis regulator *SARD1* (Q9C9T2, AT1G73805), peroxidase N1 (Q9XIV8, AT5G64120), the central immune coreceptor *SOBIR1* (Q9SKB2, AT2G31880) and the siRNA-related RNA polymerase *RDR6* (Q9SG02, AT3G49500). In *NOV* knockdown plants, these defense-associated genes were expressed at levels similar to, or lower than the baseline expression levels of mock-infected plants (Figure 1I). We interpret these results as an indication that *NOV* is required for the full transcriptional response to TRV infection, with a disproportionate effect on defense-associated genes.

### NOV contains eukaryotic-specific conserved structural motifs in its N-terminus

Beyond the C-terminal half of NOV that includes the Tiamat domains, plant NOV proteins contain an N-terminal region of ∼1000 amino acids that is absent from bacterial Tiamat proteins. This region contains nuclear localization signals, multiple predicted high-disorder stretches, and alpha-helical structured regions. To characterize the conserved content of this N-terminal region, we manually curated plant NOV homologs spanning major plant clades (Supplementary Table 3), aligned them, and partitioned the alignment into conserved regions corresponding to discrete structural- and sequence-similarity units. This analysis identified three conserved structural motifs in the N-terminal region of NOV, separated by stretches predicted to be mostly disordered regions (Figure 2A-B). These structural motifs are functionally unannotated, and belong to uncharacterized structural clusters in the AlphaFold cluster database (Barrio-Hernandez et al., 2023). However, the *nov-1* mutation previously described as affecting *A. thaliana* vascular development, Pro855>Ser (Tsugeki et al., 2009), is changing a highly conserved residue in the third of these conserved motifs, supporting the N-terminal region’s functional importance (Supplementary Figure 4).

**Figure 2:**
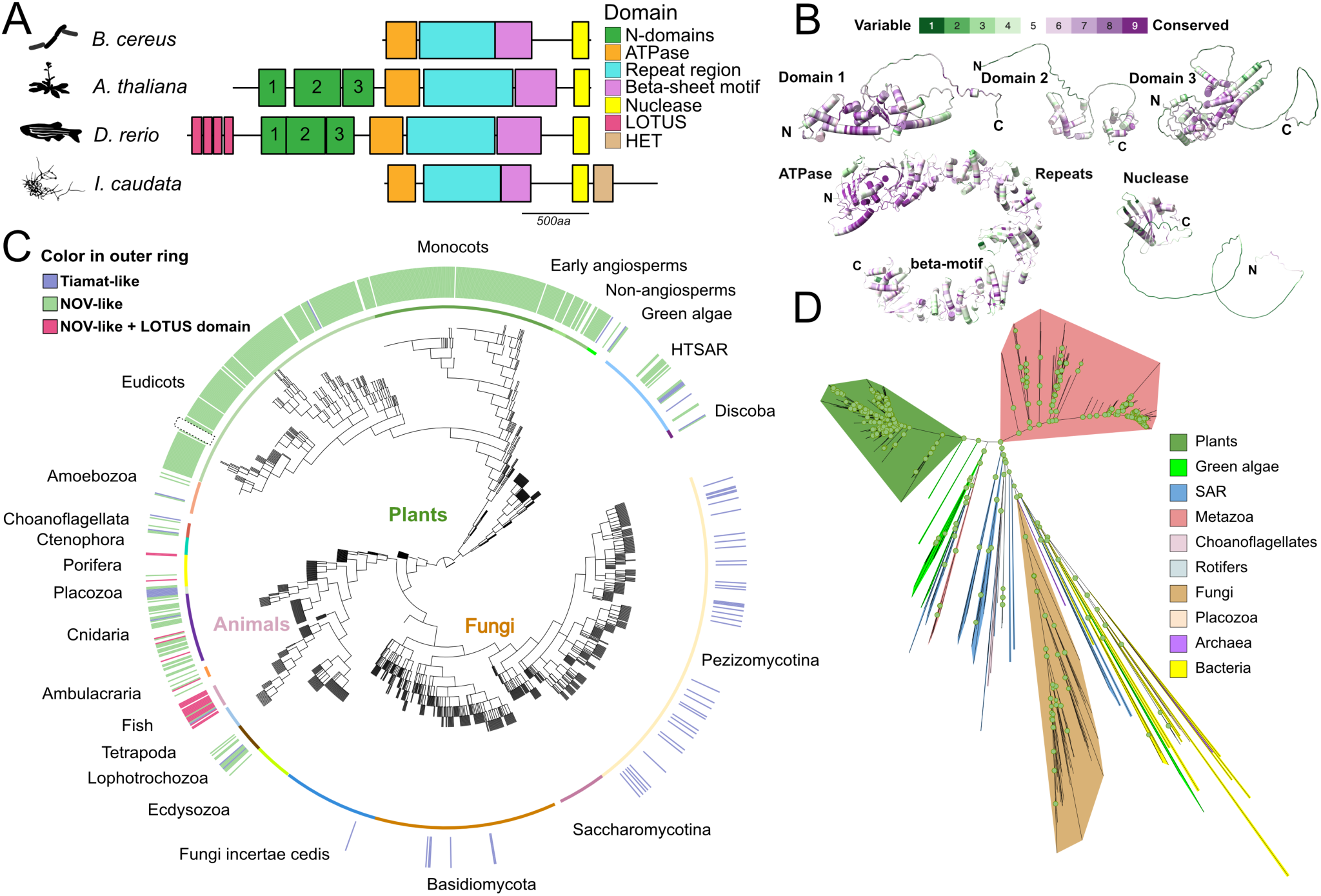
NOV homologs are widespread in plants and animals. **A** NOV consists of a Tiamat-like C-terminal half, containing the ATPase (orange), solenoid repeats (cyan), a beta-sheet motif surrounded by helical repeats (purple) and NOV_C nuclease domain (yellow). Eukaryotic NOV proteins carry an extended N-terminal region which is conserved across plant and animal homologs. This N-terminal region contains 3 unannotated conserved structural motifs (green). Fish carry additional repeated LOTUS domains (pink). Some fungal Tiamat-like homologs carry a heterokaryon incompatibility (HET) domain (brown), but are missing the N-terminal region. Species and proteins shown: *Bacillus cereus FORC_005* TmtA Refseq WP_046392403, *Arabidopsis thaliana* NOV AT4G13750, *Danio rerio* wu:fj29h11 UniprotID A0A0R4IS14, *Immersiella caudata* UniprotID A0AA39THY1. **B** The structures of the conserved NOV domains in eukaryotes are colored according to ConSurf conservation score (Yariv et al., 2023) (green-low to purple-high) projected onto the AtNOV AlphaFold3 predicted structure (see Methods). **C** NOV homologs are nearly universally present across plants, and prevalent in multiple animal lineages. No full NOV homologs were detected in Ctenophora, Ecdysozoa and Tetrapoda. In fungi, Tiamat-like homologs lack the N-terminal domains and are sparsely detected. The outer ring represents the presence of proteins containing the 3 Tiamat domains in addition to the eukaryotic-specific N-terminal motifs in green, or only the 3 regions found in Tiamat in blue. Animal proteins containing at least one N-terminal LOTUS domain in addition to the seven regions are marked in pink. The inner ring is colored according to the named taxa, Non-angiosperms include Bryophytes, Polypodiopsida, lycophytes and a gymnosperm. The phylogenetic tree was constructed from a curated

NOV was previously described as a plant-specific protein (Tsugeki et al., 2009). The identified homology of NOV to a bacterial protein suggested a more widespread presence across the tree of life. We therefore generated HMM profiles from the seven domains we defined (Figure 2A-B). The domains used were the three structural motifs in the N-terminal region, the ATPase domain, the alpha-solenoid repeat region, a conserved beta-sheet motif surrounded by helical repeats, and the NOV_C nuclease domain (Figure 2A-B). We then applied the seven HMM profiles, built from the aforementioned protein domains of plant NOVs, iteratively against a taxonomically balanced UniProt reference proteome dataset (Supplementary Table 4), extracting matching regions of proteins with hits to all seven profiles, realigning, and rebuilding the profiles until convergence at three rounds. We then used the final profiles to search a curated database for eukaryotic sampling, including proteomes from Phytozome, LukProt and Uniprot (see Methods, Supplementary Table 4), and classified hits by domain composition (Supplementary table 4, Figure 2C).

### NOV is broadly conserved in plants but pseudogenized in *Thlaspi arvense*

NOV is nearly universally conserved across land plants, consistent with its identification as an essential gene for early embryogenesis (Meinke, 2020; Tsugeki et al., 2009). We identified NOV homologs in bryophytes and ferns, indicating its presence in the common ancestor of land plants (Figure 2C). Notably, despite its broad conservation across angiosperms in general and Brassicaceae specifically (Figure 2C), *NOV* appears to be a pseudogene in *Thlaspi arvense*, a close relative of *A. thaliana*. Across multiple *T. arvense* accessions (Bird et al., 2026), the *NOV* gene has premature stop codons and disrupted gene structure relative to the intact *A. thaliana* gene (Supplementary Figure 5). No other intact copies of *NOV* were identified in the full genome assemblies of *T. arvense*. Analysis of published RNA sequencing reads (Bird et al., 2026) identified no transcription of the *NOV* pseudogene in 6 out of the 7 *T. arvense* accessions, while the transcripts in accession Ames32873 are coding for frameshifted open reading frames. This indicates that *NOV* is not strictly essential in all Brassicaceae genera, and that its essential functions in *A. thaliana* are compensated by unknown factors, or not required for survival in *T. arvense*.

### NOV/Tiamat proteins in animals and fungi

Beyond plants, our searches identified NOV homologs across multiple animal lineages. Animal NOV homologs contain all of the plant NOV domains, including the N-terminal region (Figure 2A). Certain animal homologs, particularly in fish, carry at the N-terminus one or more LOTUS domains, a domain associated with RNA binding (Figure 2A,C) (Ding et al., 2020). The presence of full NOV homologs in both plants and animals suggests that NOV emergence predates the split between these lineages.

We did not detect NOV homologs in several animal lineages, including Ecdysozoa, the group containing Arthropods and Nematodes. Although NOV is present in cartilaginous, bony and lobe-finned fishes (Chondrichthyes, Actinopterygii and Sarcopterygii) it was not detected in Tetrapods. This is in contrast to the ubiquitous conservation of NOV in all land plant lineages, and indicates a separate evolutionary trajectory of animal NOVs.

In contrast to NOV’s widespread presence in animals and plants, we did not detect the full NOV domain architecture in fungi. To investigate the relationship between bacterial

Tiamat, fungal homologs, and plant/animal NOVs, we used the Tiamat HMM profiles of DefenseFinder to identify all Tiamat-like proteins in the balanced Uniprot dataset, including both prokaryotes and eukaryotes, and filtered for proteins containing both the ATPase and NOV_C domains (Supplementary Table 4). Despite close evolutionary relationship of fungi to animals as sister clades of opisthokonts, fungal homologs were observed to lack the N-terminal region and contain only the Tiamat-like core, with some homologs additionally carrying heterokaryon incompatibility domains (Figure 2A, Figure 2C, Supplementary Table 4). We generated a phylogenetic tree based on the protein region spanning from the start of the ATPase domain to the end of the NOV_C nuclease domain (Figure 2D). In this tree, fungal Tiamat-like homologs are well separated from both animal and plant NOVs. Together with the absence of the N-terminal region in these proteins, these observations are consistent with independent horizontal transfers of Tiamat-like genes from prokaryotes into fungi, which have well-documented propensity for such transfers (Richards et al., 2011). However, we cannot exclude ancient vertical inheritance from the ancestor of opisthokonts followed by extensive divergence and independent loss from multiple fungal lineages.

Eukaryotic genome database (see Methods, Supplementary Table 4). Dotted box highlights absence of NOV in *Thlaspi arvense* accessions (Supplementary Figure 5). **D** Phylogenetic tree of Tiamat homologs, based on MAFFT alignment of the sequence from the start of the ATPase domain to the end of the NOV_C domain (Supplementary Table 4). The tree was constructed using IQ-TREE 3 (Wong et al., 2026), Well supported nodes (SH-aLRT & UFBoot > 80) are marked with green dots.

## Discussion

This work identifies plant NOV proteins as related to the bacterial antiviral protein Tiamat, and discovers widespread animal homologs of NOV. Tiamat and eukaryotic NOV proteins share a core composition over more than 1000 amino acids that spans multiple protein domains (Figure 1A, Supplementary Figure 1, Figure 2A-B). We hypothesized that this protein’s similarity to a bacterial immune protein could reflect shared functions in anti-viral immunity. In agreement, *NOV* knockdown increases PVX accumulation *in planta* and dampened immunity-related gene expression in response to TRV (Figure 1). Together, these results reveal NOV as a conserved eukaryotic protein with functions in plant development (Tsugeki et al., 2009) and immunity, sharing domain composition with a bacterial antiviral protein.

The transcriptional effects of *NOV* knockdown point towards impaired induction of immunity genes in response to TRV infection. As the TRV transcriptome was sampled at the timepoint equivalent to the inoculation of a secondary *Agrobacterium*-induced PVX infection, it is possible that the increased PVX accumulation is a result of the immunocompromised state of *NOV* knockdown plants. We note that while PVX accumulation was significantly increased in *NOV* knockdown plants, AbMV and CpCDV infection outcomes were unaffected. This suggests differential sensitivity of different viruses to *NOV*-related transcriptional changes or a specific role for NOV in PVX infection.

Impaired immunity-related gene induction was previously reported for mutants in plant transcriptional and RNA-processing machinery proteins. In *A. thaliana*, mutants in the Mediator-complex tail subunits MED14, MED15 and MED16 show reduced induction of defense-associated genes across multiple pathogen and hormone stimuli, while the induction of other genes was not affected (Wang et al., 2016; Wathugala et al., 2012; S. Zhang et al., 2026; X. Zhang et al., 2013). Mutations in Elongator subunit 2 cause a delay in immunity-related gene expression (DeFraia et al., 2010). NOV itself is a nuclear protein (Tsugeki et al., 2009), and we hypothesize that its roles in immunity and development can be related to functions in fundamental nuclear processes that are essential for the execution of certain transcriptional programs.

NOV is broadly conserved in plants and animals, indicating that the eukaryotic version of this protein, including the N-terminal domains, originated before the split between plants and opisthokonts. This is supported by both the conserved domain composition (Figure 2A) and the phylogenetic analysis of the conserved Tiamat-like region (Figure 2D). The functions of animal homologs of NOV are yet largely unknown. An interesting connection between NOV proteins, stress and immunity in non-plant systems comes from the observation that a NOV homolog is the top gene enriched in the dinoflagellate symbiont *Cladocopium goreaui* of healthy coral tissue, when compared to corals with stony-coral tissue loss disease (Beavers, 2024).

NOV and Tiamat proteins are members of the larger, previously defined protein family of Sacsin-repeat containing proteins (Anderson et al., 2010). This family is defined by the Sacsin Repeat supradomain, which includes the ATPase domain and the solenoid repeat section (Perna et al., 2022). In *A. thaliana,* we inspected two additional uncharacterized members of this superfamily that were previously indexed (Anderson et al., 2010). AT3G48770 has an N-terminal BED-type zinc-finger domain, a single sacsin repeat, and is terminated by the beta-sheet motif identified in this study (Supplementary Figure 6A). AT5G23110 shares human Sacsin’s organization of three sacsin repeats followed by one copy of the beta-sheet motif, but ends in a C-terminal C3HC4 zinc-finger domain instead of Sacsins’s HEPN and J-domains (Supplementary Figure 6B-C). Sacsin-repeat containing proteins are associated with variable nucleic acid interacting domains - NOV and Tiamat carry a PD(D/E)XK nuclease, fish NOV homologs additionally carry a putative RNA-interacting LOTUS domain, Sacsin carries an RNA-binding HEPN domain (Anderson et al., 2010; Longo et al., 2025), and the abovementioned two *A. thaliana* proteins carry zinc finger domains. The co-occurence of nucleic-acid related domains with sacsin-repeat containing proteins suggests that these proteins are frequently involved in nucleic acid interaction. As NOV is a nuclear protein (Tsugeki et al., 2009) and we observe that perturbation of *NOV* affects the robustness of a specific transcriptional response, this suggests that the Sacsin family may be involved in regulatory RNA-related processes.

Taken together, our results expand the known homologies between bacterial immune systems and eukaryotic proteins, and especially the small set of ancestral immune modules currently characterized in plants. We demonstrate that such homology can serve as a guide to identify new plant proteins involved in immunity. The homology between plant and animal NOV proteins and bacterial Tiamat could assist future studies, as it opens experimental avenues to study different aspects in different systems. Beyond that, NOV serves as an example of an ancient bacterial immune protein that became embedded in essential processes in plants.

## Methods

### DefenseFinder search in plants

To search for defense system signatures in *Arabidopsis thaliana*, we used the DefenseFinder tool version 3.1.0 (Tesson et al., 2022) with default settings against the Araport11 proteome (Supplementary table 1A-B).

### HMM profile definition of NOV domains

We manually curated 97 plant homologs of NOV (Supplementary table 3), covering most major clades, and aligned them by MAFFT. The alignment was partitioned into 7 regions corresponding to conserved sequence and structural elements, and a separate HMM profile was built for each region. These profiles were applied in an iterative HMMsearch against a taxonomically balanced UniProt dataset where 6 representatives from each family were selected across eukaryotes and prokaryotes, downloaded using a publicly available API tool (download_reference_proteomes.py from github.com/nickatirwin/phylogenomic_scripts). For each iteration, we extracted regions matching the HMM profiles from proteins containing all 7 conserved regions - classified as NOV - realigned them, and rebuilt new profiles. Iteration continued until no new hits were recovered, which occurred after 3 rounds. No prokaryotic proteins were found to contain any of the N-terminal NOV domains.

To visualize the presence/absence of NOV and Tiamat across eukaryotes, we used the LukProt dataset (an extension of EukProt) as the phylogenetic backbone, since it provides broader sampling across eukaryotes (Richter et al., 2022; Sobala, 2024). Genomes with contamination above 5% and/or BUSCO completeness below 60% were excluded. To broaden the sampling of fungi and plants, we replaced the fungal and Chloroplastida clades of the LukProt tree with denser datasets: for fungi, we used the UniProt reference proteome set described above; for plants, we used Phytozome, which provides a larger and more uniformly assembled set of plant genomes as well as pangenomic assemblies. We applied the third-round HMM profiles to all three datasets (LukProt, fungal UniProt subset, plants) to identify NOV- and Tiamat-like homologs, then assembled a chimeric phylogeny by grafting NCBI common-tree topologies for fungi and plants retrieved via ete3 onto the LukProt backbone at the positions of the original fungal and Chloroplastida clades. Plant pangenomic datasets were grafted to their common ancestor based on NCBI taxonomy. Because this phylogeny is chimeric and not based on sequence distances, this tree was only used for presence/absence visualization across clades.

### Detection of NOV in Thlaspi arvense

To investigate absence of NOV protein homologs in *Thlaspi arvense*, genome assemblies for 7 different accessions from the PennyPan *T. arvense* pangenome project (AK34W, LorettoMN, MN106, Tibet33, Ames 32873, MN134 and PI650286 (Bird et al., 2026)) and *Schrenkiella parvula* as a closely related outgroup (Nikolov et al., 2019) were downloaded from phytozome-next.jgi.doe.org in May 2026 (Goodstein et al., 2012). A BLASTn v2.17.0 search was performed on the genomic sequences of the *T. arvense* accessions to identify homologous regions corresponding to the coding sequence of *NOV* from *A. thaliana* and *S. parvula NOV* (Sp7g11520), with the settings dc-megablast and an E-value threshold of 1e-5. Only one *NOV* locus per genome was identified. The genomic regions corresponding to the *NOV* locus were extracted using SAMtools faidx with 30kb flanking windows (Danecek et al., 2021). To define the exon structure of *NOV* gene models in *T. arvense,* Exonerate v2.4.0 (Slater & Birney, 2005) was used with the protein2genome model to search these regions for homology to the *A. thaliana* NOV protein sequence, reporting up to ten alignments per query (--bestn 10) so that decayed gene regions split into several alignments were fully captured. This was also done using *S. parvula* NOV as a more closely related species, and the same results were observed. Disabling mutations were recorded: in-frame premature stop codons, frameshifts, and splice sites departing from GT/GC donor or AG acceptor dinucleotides. To compare accessions and species, exons and mutations were visualized on the Araport11 *NOV* gene model by their position in the *At*NOV protein. The in-frame part of each gene was defined as the stretch from the aligned start to the first premature stop codon or frameshift. Feature coordinates were plotted with gggenomes v1.2.0 (Hackl et al., 2024) in R (v4.4.0 (R Core Team, 2026)) Locus status was classified as intact (no premature stops in the exonerate model) or pseudogenized (one or more premature stops). The Ames32873 region was displayed in reverse-complement orientation to normalise for an assembly-level inversion of chromosome 7 (Bird et al., 2026).

### Structural prediction and alignments

We used AlphaFold3 (Abramson et al., 2024) with default settings and random seeds to predict the protein structures of *Bacillus cereus* FORC005 Tiamat (Uniparc: UPI00061DE4E5, RefSeq: WP_046392403), *A. thaliana* and *N. benthamiana* NOV (AT4G13750, NbL01g24040), an *Immersiella caudata Tiamat*-like protein (Uniprot: A0AA39THY1), *Danio rerio* wu:fj29h11 (Uniprot: A0A0R4IS14) AT5G23110 (Uniprot: Q9FN44) and *Homo sapiens* Sacsin (Uniprot: Q9NZJ4). The predicted structure of AT3G48770 (Uniprot: Q9M305) was downloaded from AlphaFoldDB under the accession AF-Q9M305-F1. The AF3 predicted structures were inspected in ChimeraX v1.10 (Meng et al., 2023) to confirm consistent model prediction.

TMalign version 20240303 (C. Zhang et al., 2022, 2026; C. Zhang & Pyle, 2022; Y. Zhang & Skolnick, 2005) was used to assess the structural similarity between segments of Tiamat and AtNOV predicted structures, spanning the ATPase to the nuclease domain. The structures were divided into 5 segments based on visual inspection and common folds and the segments were aligned pairwise to each other. Structures and protein alignments were imported into ChimeraX (Meng et al., 2023) and conserved catalytic residues in the ATPase and nuclease domains were highlighted (Millman et al., 2022; Steczkiewicz et al., 2012).

EMERALD-UI (Grigorjew et al., 2023; Preoteasa et al., 2026) was used to align windows of Tiamat WP_046392403 and AtNOV^1156-2729^. This region of AtNOV was used as the 1159-2709 Tiamat HMM aligned region truncates 3 amino acids of an alpha-helix motif, and also includes the last 20 amino acids following the nuclease domain. The entire region spans the ATPase domain to the nuclease domain, and alignment was run using the default settings at α=0.75 and Δ=8.

### Conservation of NOV domains using ConSurf

Regions spanning the first amino acid corresponding to the HMM region of the first NOV fold to the first amino acid of the ATPase domain were extracted from proteins classified as NOV in Figure 2C, including plant and animal proteins. These 596 total proteins were clustered using DIAMOND v2.1.24 (Buchfink et al., 2021) at 90% coverage and 90% identity, down to 169 total sequences and aligned using MUSCLE v3.8.1551 (Edgar, 2004). This multiple sequence alignment was uploaded to the ConSurf colab notebook (https://github.com/BenTalLab/consurf) (Yariv et al., 2023) along with the AlphaFold3 predicted structure of AtNOV, and results were imported into ChimeraX-1.10 (Meng et al., 2023) for projecting the conserved residues onto the structure. The same was done for the C-terminal half of NOV proteins - corresponding to the first amino acid of the ATPase domain to the last amino acid of the NOV_C domain as determined through profile HMM matching.

### Phylogenetic reconstruction of NOV/Tiamat homologous region

We used the Tiamat profile-HMMs from DefenseFinder (Tesson et al., 2022) in an HMMsearch using HMMER v3.4 (Eddy, 2011) to identify additional Tiamat homologs across bacteria, archaea and eukaryotes from the UniProt dataset in Supplementary table 4. Protein domains were annotated using the seven NOV HMM profiles and sequences containing the ATPase and NOV_C domains were kept. The regions spanning these two HMM profiles were extracted and aligned using MAFFT (Katoh & Standley, 2013), gaps were trimmed using TrimAl (Capella-Gutiérrez et al., 2009) and a phylogenetic tree was reconstructed using IQ-TREE 3 (Wong et al., 2026) using the Q.PFAM+I+G4 model which was inferred from modelfinder. Branch support was assessed using Sh-ALRT + UFBoot with 1000 replicates. The tree was imported into iTOL, rooted at midpoint and clades were automatically annotated based on the embedded NCBI identifiers in the protein IDs and manually colored based on taxonomic groups.

### TRV-VIGS knockdown in *N. benthamiana*

We restriction cloned fragments of the *NbNOV* CDS (NbL01g24040), the NLR gene *NbPrf* (NbL08g09580) – an immune receptor specific for a *Phytophthora* effector, not known to be involved in viral immunity (Kim et al., 2002), or the *E. coli GUS* gene from pENTR-GUS into the TRV2 cloning vector pYL156 (Liu et al., 2002) (addgene #148969) and introduced plasmids to *Agrobacterium tumefaciens* GV3101 through electroporation. *A. tumefaciens* clones carrying TRV1 (pYL192 - addgene #148968) and TRV2 silencing constructs were both adjusted to OD 0.1, mixed 1:1 in standard infiltration buffer (10mM MES pH 5.7, 10mM MgCl_2_ and 150µM acetosyringone) and infiltrated into the first true leaf of 2 week old *N. benthamiana* seedlings. For mock infections, an agrobacterium clone carrying the pH7WGF2 empty GFP construct was used to match the total agrobacterium load in the TRV treatments. Shoot apexes, including non-expanded leaves, were harvested after 10 days for RNA extraction. Total nucleic acids were extracted using guanidine hydrochloride buffer and 96-well spin columns as previously described (Leite Montalvão et al., 2026), and RNA was isolated by treating samples with DNAse I. Gene expression levels were measured by RT-qPCR using RevertAid RT kit with random hexamer primers (Thermo Fisher) for reverse transcription and LUNA (NEB) for qPCR, primers for *Nb*eIF4A (NbL18g11970) were used as an endogenous gene control. Primers for qPCR were made 3’ downstream of the VIGS target region. Primers and sequences and the lengths of cloning inserts are listed in Supplementary Table 5. Plants were grown in a growth chamber with 16h full spectrum light, 8h darkness at 24°C and 65% relative humidity.

### PVX, CpCDV and AbMV infection assays

Infectious agrobacterium clones carrying genomes of PVX, CpCDV or AbMV, were mixed into standard infiltration buffer (10mM MES pH 5.7, 10mM MgCl_2_ and 150µM acetosyringone) at OD0.05 for PVX, OD 0.1 for CpCDV, and OD 0.1 for the bipartite geminivirus AbMV A and B of which the two *Agrobacterium* clones carrying each part were mixed 1:1. Plants previously infiltrated with TRV VIGS constructs or mock infected were infiltrated with viral clones 10 days after initial TRV infiltration. Infiltration was performed as a 2cm diameter circle into the top fully expanded leaf of the plants. Plants were grown for an additional 10 days, apexes were harvested and total nucleic acid was isolated as described above. For PVX and TRV, we isolated RNA, normalized it to 50ng/µl where 500ng was used for cDNA synthesis and subsequent RT-qPCR on 1µl of 1/10 diluted cDNA using *Nb*eIF4A primers as internal control and PVX coat protein primers as a measure for viral load. Tobacco rattle virus RNA2 was quantified through RT-qPCR targeting the coat protein RNA. For geminiviruses we performed DNA qPCR on extracted DNA at 10ng/µl using NbNOV as a single copy gene control (verified by blastn of the gene to the NbLAB360 genome (Ranawaka et al., 2023)) and coat protein gene primers corresponding to the two different viruses for viral load.

Statistical analysis was performed in R (v4.4.0 (R Core Team, 2026)) using the R library lme4 (Bates et al., 2015) on *Δ*Ct values. *Δ*Ct values were calculated through the difference of mean Ct values from qPCR technical triplicates between the Ct value of loading control (*Nb*eIF4a for RNA samples, *NbNOV* for DNA samples) and the Ct value of the amplicon of interest. Independent experimental replicates were used as a random error factor - with the linear model design *Δ*Ct ∼ infection treatment + (1|experimental replicate). P-values were adjusted using Benjamini-Hochberg false discovery rate correction. For visualization, figures (box and bar plots) were presented with the linearly transformed *ΔΔ*Ct values as the normalized values on the y axis, using TRV-GUS as the control treatment for each independent experimental batch to calculate the *ΔΔ*Ct values by 2^-(mean *Δ*Ct_GUS per batch - *Δ*Ct_test).

### RNA sequencing

Samples for RNA sequencing were chosen from two experimental batches where each experiment included three treatments: mock (no TRV), TRV-GUS (silencing control), and TRV-NOV, with four plants per treatment per batch. Samples were selected to represent the distribution of *NOV* expression and TRV load based on prior RT-qPCR data. Extracted RNA was sent for poly-A-enriched library preparation and sequencing to Genewiz Germany GmbH (Leipzig, Germany). Sequencing was performed on an Illumina NovaseqX platform using 150bp paired end sequencing aimed at 40M total reads per sample. We used the NF-core RNAseq version 3.14 pipeline (Patel et al., 2026) for read alignment to the *N. benthamiana* LAB360 genome assembly (Ranawaka et al., 2023). The region corresponding to the TRV-Nb*NOV*-1 silencing construct in the *N. benthamiana* genome was hardmasked with N to prevent VIGS reads from inflating endogenous *Nb*NOV quantification, and sequences corresponding to the TRV genome were appended to this genome .fasta and .gff for read mapping to the viral genome. Within the pipeline, reads were adapter- and quality-trimmed with Trim Galore, aligned with STAR, and quantified with Salmon in the star_salmon mode; gene-level count matrices were generated by the pipeline’s tximport step. Library strandedness was set to automatic detection and inferred as unstranded (RSeQC infer_experiment, ∼50:50 sense:antisense). Raw viral reads were quantified using samtools coverage (Danecek et al., 2021) while TPM values for the TRV genes (16k, RdRp, and MP in RNA1; and Cp, 2b and inserts corresponding to the silencing constructs in RNA2), as output by Salmon were used for data analysis.

One TRV-NOV-1 sample from TRV-only Rep1 failed pre-sequencing RNA quality control, leaving three biological replicates in this batch. One mock sample was excluded from the TRV-experiment analysis after detection of thousands of TRV RNA1 reads, though no TRV RNA2 reads were detected, indicative of unspecific contamination. Remaining mock samples contained at most a few dozen uniquely mapped TRV1 reads at shallow, dispersed coverage, a sign of low-level index hopping from co-sequenced infected samples (Supplementary table 2) and were retained. No cross-contamination between the two TRV2 constructs were detected, by counting reads mapping to the GUS/NOV inserts.

DESeq2 v1.46.0 (Love et al., 2014) was used for differential gene expression analysis using R Statistical Software (v4.4.0; (R Core Team, 2026)), with the gene-level count matrices used as input. Viral-contig features were removed prior to host differential-expression analysis. Multifactorial design accounting for batch effects was used for DESeq2 analysis. Pairwise contrasts (TRV-NOV vs TRV-GUS; TRV-GUS vs mock; TRV-NOV vs Mock) were extracted, and log2 fold-change estimates were shrunk with the ashr estimator (Stephens, 2017). Genes with a Benjamini–Hochberg-adjusted p < 0.05 and |log2FC| > 0.4 were considered statistically significant.

To assess batch effects and sample variance, gene counts were transformed by variance stabilizing transformation and normalized to library size to yield approximate homoskedastic values in DESeq2 (Love et al., 2014), which was used to calculate principal component variation using the DESeq2 plotPCA function.

#### Functional annotation of Nicotiana benthamiana LAB360 gene models

As the NbLab360 genome annotation provides limited functional information, NbLab360 protein models (CDS-derived amino-acid sequences) were assigned annotations by homology search with DIAMOND (v2.1.24); (Buchfink et al., 2021) in blastp mode against two reference sets: Arabidopsis thaliana proteins (Araport11/TAIR) and UniProt Swiss-Prot reviewed proteins (Viridiplantae). Hits were retained at ≥30% identity, ≥50% query coverage, and E-value ≤ 1×10^−10^ for both, keeping the single best hit per query if higher than 40% identity for annotations, with preference for TAIR IDs in cases of ties, and defaulting to NbLAB360 identifiers when hits did not qualify (i.e <30% identity). The resulting TAIR or Uniprot homolog was used for gene annotation in figures 1 F-H and Supplementary Figure 3B. Gene Ontology (GO) annotation was determined through org.At.tair.db based on TAIR identifiers for genes where such a homolog was detected. For defining Defense-related and heat-stress related gene sets, GO:0006952 (Defense response) and GO:0009408 (Heat response) were used. In overlapping cases, ‘Defense response’ was preferred over ‘Heat response’.

Transposable-element annotation of the NbLab360 gene models was assessed by DIAMOND blastp of the amino-acid models against the REXdb Viridiplantae TE protein database v4.0; (Neumann et al., 2019), retaining the best hit per gene at ≥30% identity, ≥30% query coverage, and E-value ≤ 1×10^−5^. TE hits were joined to the REXdb classification table to assign each TE-associated gene a hierarchical classification (class, order, superfamily, and lineage). Genes with a qualifying REXdb hit were flagged as TE-associated for downstream analyses. Genes containing “transposon” in their uniprot or TAIR descriptions were also flagged as TE-associated.

## Acknowledgements

We thank Detlef Weigel for supporting this work and for his advice and mentorship. We thank Clémence Marchal for sharing Potato Virus X clones, and Chaonan Shi, Yu Zhou and Huang Tan for experimental advice. We thank Julian Pischos for his assistance with experiments. We thank Naja Faerber, Lukas Fiedler and Adi Millman for their helpful comments on the draft manuscript. We thank Kevin A. Bird for his advice on Thlaspi genome analysis. We thank Aude Bernheim for her advice. The study was funded by Max Planck Society funding to Detlef Weigel, EMBO Long Term Fellowship (ALTF 851-2021) and HFSP Long Term Fellowship (LT 0044/2022-L) to Gal Ofir.

## Data availability

Raw RNAseq reads can be found under the ENA study accession PRJEB127217.

## Use of LLM tools

Claude (Sonnet and Opus) by Anthropic was used for assistance with textual editing, proofreading and code. All code was manually reviewed before use, and run without agentic use.

## Supplementary material

Supplementary Table 1 - DefenseFinder hits **A** Complete systems **B** Individual HMM profile hits to *A. thaliana* genes

Supplementary Table 2 - DEGs in N. benthamiana TRV-NOV silenced plants **A** GUS vs Mock, **B** NOV vs Mock, **C** NOV vs GUS, **D** Comparison table of A&B **E** TRV TPM values **F** TRV contig counts RNAseq

Supplementary Table 3 - Plant NOV proteins used to generate HMM profiles of the 7 domains, with domain coordinates on each

Supplementary Table 4 - Proteomes used to search for NOV homologs in other eukaryotes from Uniprot, LukProt and Phytozome, and the hits found with domain coordinates on each. **A** Presence/absence matrix of Tiamat, NOV and LOTUS domain presence **B** NOV hmm profile hits across LukProt, fungi and plants with PFAM annotations and NOV domain locations. **C** Tiamat HMM profile hits across UniProt reference proteome dataset **D** NOV domain profile HMM hits on proteins from C. **E** Proteins containing NOV ATPase + NOV_C domain profile hits from D. **F** Protein coordinates from proteins in E used for phylogenetic tree **G** Lookup table for A-B containing a list of the proteomes used **H** LukProt genomes used, prefiltered with BUSCO >60% and OMark contamination <5% (from original LukProt dataset calculations). **I** Plant genomes used **J** Fungal genomes used **K** UniProt dataset used

Supplementary Table 5 - Primers for PCR (**A)** and sequences used (**B**)

**Supplementary Figure 1:**
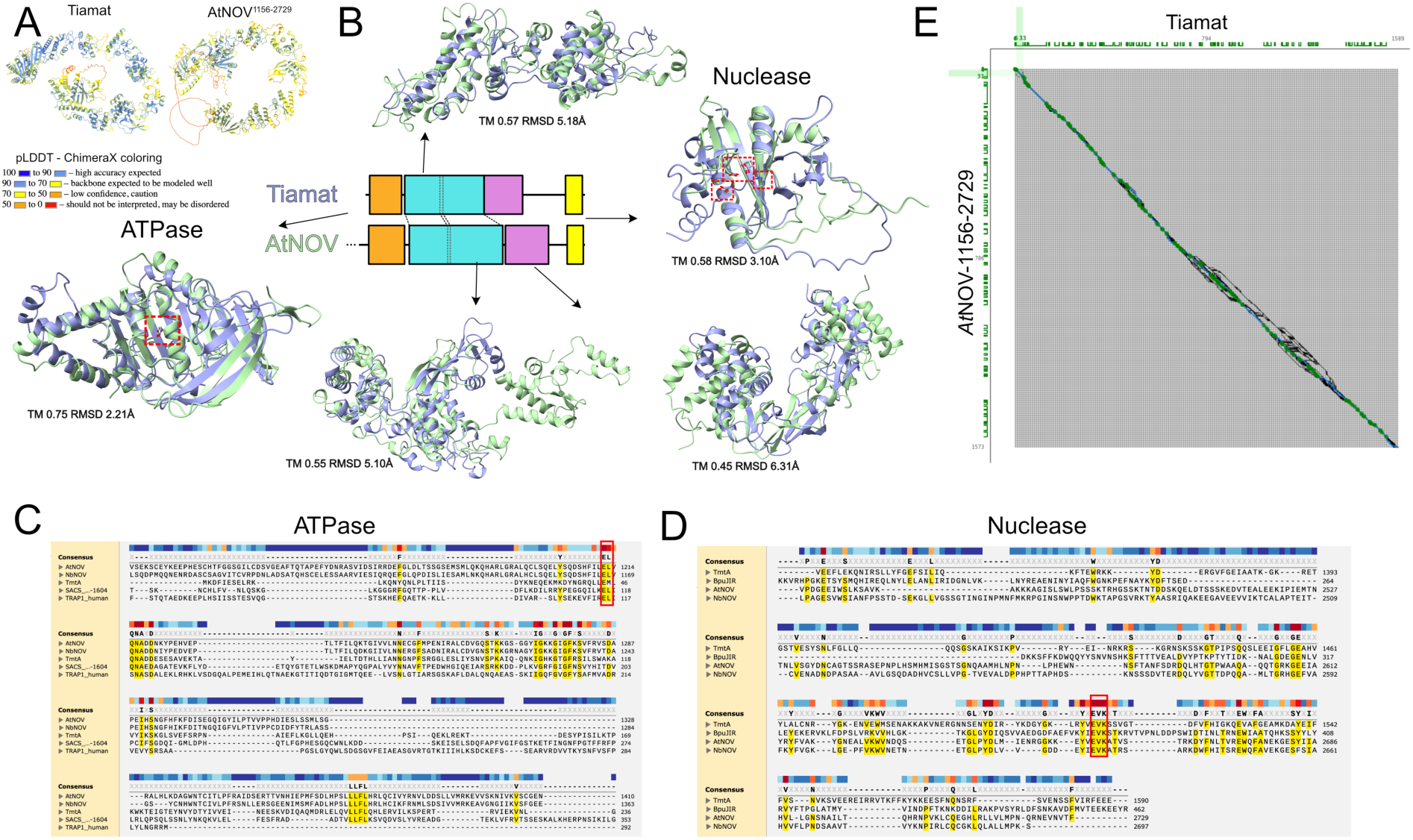
Structural alignments of AtNOV and Tiamat. **A** AlphaFold3 predicted structures of Tiamat and the alignable section of AtNOV, colored by prediction confidence pLDDT scores according to the ChimeraX coloring scheme. **B** Structural alignments of individual Tiamat (Blue) and NOV (Green) regions with TMalign TM and RMSD values to the AtNOV aligned region. Regions aligned are shown on the Tiamat and NOV linear representations where the ATPase domain is orange; the alpha-solenoid repeat region in cyan - split into two regions, the second of which contains an additional solenoid repeat in AtNOV; the beta-sheet domain in magenta and the nuclease domain in yellow. Dotted boxes highlight active site residues. A mutation in the ATPase active site residue (E44) abolished Tiamat’s antiviral activity (Millman et al., 2022). This residue corresponds to E1212 in AtNOV. In the nuclease domain dotted boxes highlights the canonical residues required for activity in PD(D/E)XK nucleases corresponding to in AtNOV: E2609, D2640, E2654, K2656; Tiamat: E1458, D1479, E1511, K1513. **C** Clustal sequence alignment of the ATPase domains, red box highlights the catalytic site. Sequences are of *Arabidopsis thaliana* NOV, *Nicotiana benthamiana* NOV, *Bacillus cereus* Tiamat, *Homo sapiens* SACS repeat region 1 and TRAP1 ATPase from *Homo sapiens* as an outgroup based on HMM profile match to the NOV ATPase domain. **D** Clustal sequence alignment of the PD(D/E)XK nuclease domains, red box highlight the catalytic site.Sequences are from *Bc*Tiamat, *At*NOV, *Nb*NOV, and *Bacillus pumilus* BpuJIR (A3FMN7_BACPU) as an outgroup with known enzymatic activity of this conserved domain. **E** EMERALD-UI alignment windows of Tiamat (x-axis) and AtNOV (y-axis) spanning the ATPase domain to the nuclease domain, using default settings. Green intervals indicate alignment safe windows at an α=0.75 and Δ=8. Complex regions with black lines diverging diagonally, horizontally or vertically suggest areas of high variability, and are apparent in the region of Tiamat spanning amino acids 1139-1381 between the beta-sheet motif and the nuclease domain, which is not present in NOV.

**Supplementary Figure 2:**
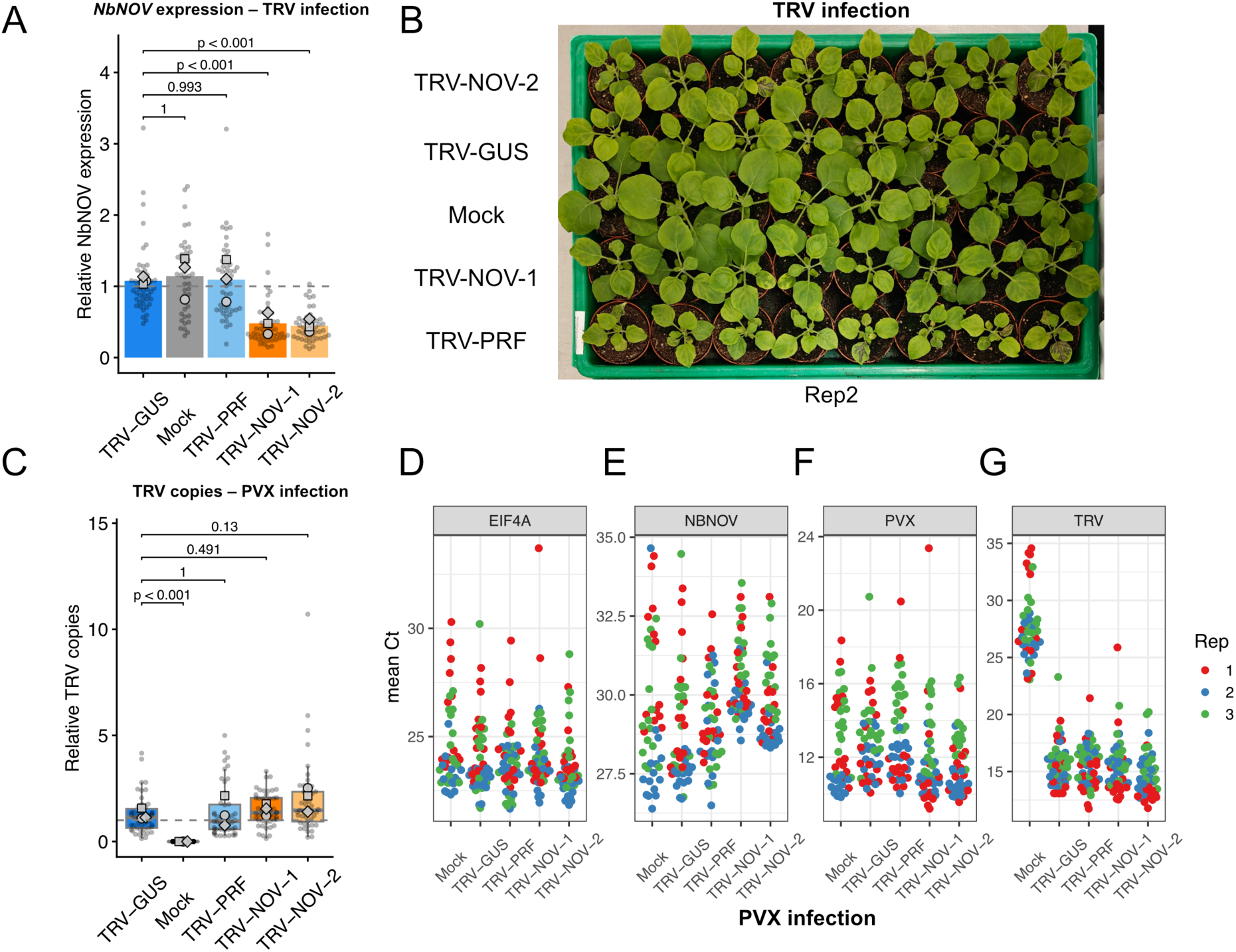
TRV-VIGS knockdown of *NbNOV*. **A** *NbNOV* is knocked down in both TRV-NOV-1 and TRV-NOV-2 treatments. Three independent experiments with 16 replicates per experiment. Means of each replicate are presented as different shapes. Relative expression normalized to TRV-GUS per replicate, dotted line is 1. Expression levels of *NbNOV* were normalized to *eIF4A*, see Methods. **B** TRV-NOV constructs do not affect plant growth compared to the control TRV-GUS infection. **C** TRV copy numbers are not different between TRV-GUS and other TRV treatments when coinfected with PVX. Data corresponds with samples from Figure 1B-C. **D** *NbeIF4A* is a stable internal control gene in samples coinfected with TRV and PVX across three experimental replicates. RNA input was normalized to 500ng for cDNA synthesis for subsequent qPCR, and qPCR was run using 1/10 diluted cDNA. Each individual point in **D-G** is a mean Ct value from raw qPCR technical triplicates, colored by replicates as in the legend. **E** Mean Ct values of *NbNOV*. **F** Mean Ct values of PVX coat protein transcripts. **G** Mean Ct values of TRV coat protein transcript.

**Supplementary Figure 3:**
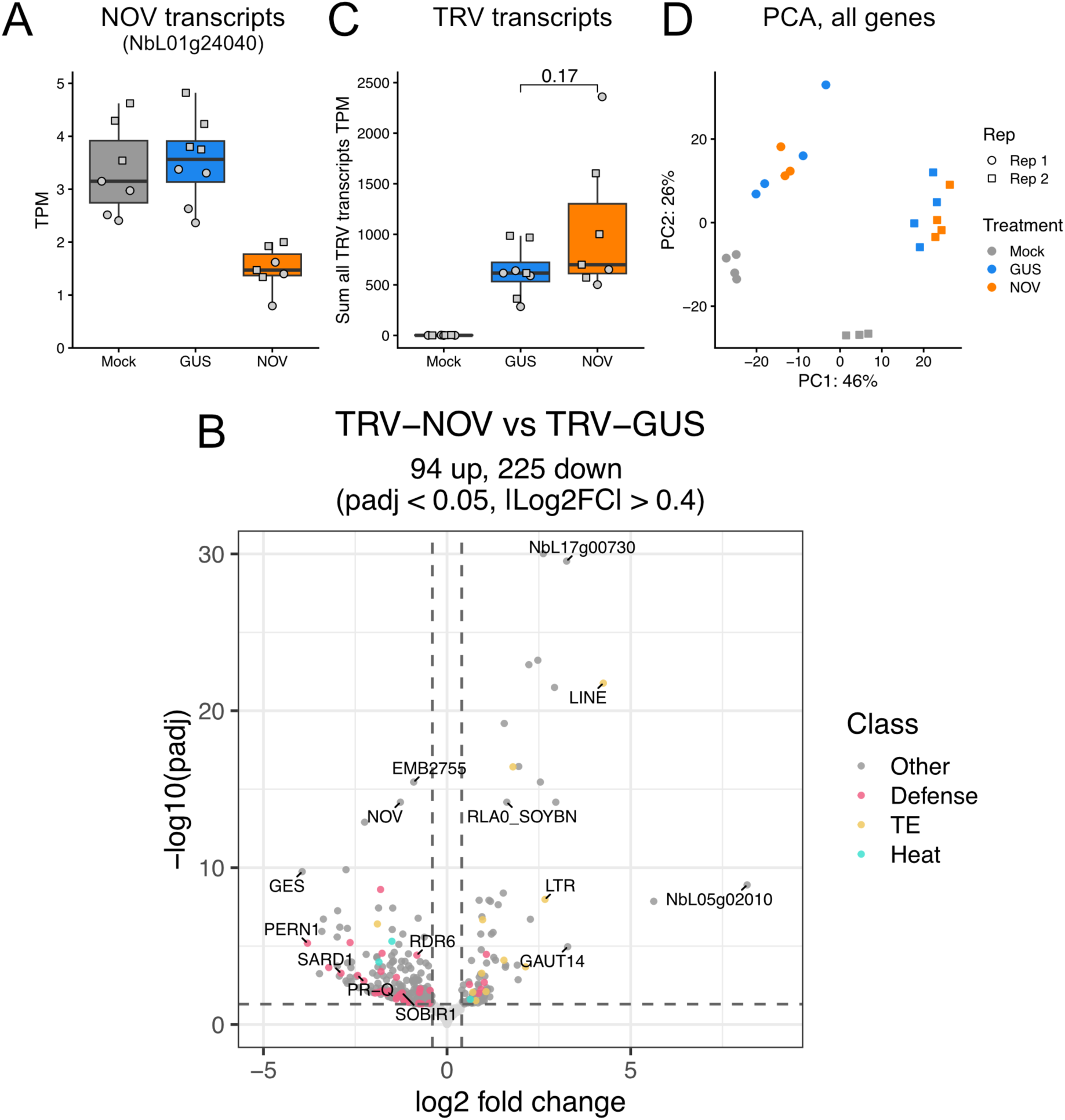
*NOV* knockdown downregulates defense genes compared to TRV-GUS control. **A** Transcript per million (TPM) of *NbNOV* (NbL01g24040) in RNAseq samples. **B** Volcano plot of differentially expressed genes between TRV-NOV and TRV-GUS treatments. Genes assigned to the GO term “Defense response” are marked in red, “Heat response” genes are marked in blue, transposable elements are marked in yellow. Specific genes with high fold change and high significance are labelled with their TAIR symbol, Uniprot symbol or NbLAB360 gene identifier. **C** Sums of TRV transcripts per million in RNAseq samples across the TRV RNA1 (16k, RdRp, and MP) and RNA2 (Cp, 2b and inserts corresponding to the silencing constructs). The p-value of a Student’s T-test is presented. **D** Principal component analysis of expression levels for all genes in RNAseq samples. Variance on x-axis is driven mainly by batch effects and y-axis variance is driven mainly by TRV presence, while intra-batch variance is low between samples containing TRV.

**Supplementary Figure 4:**
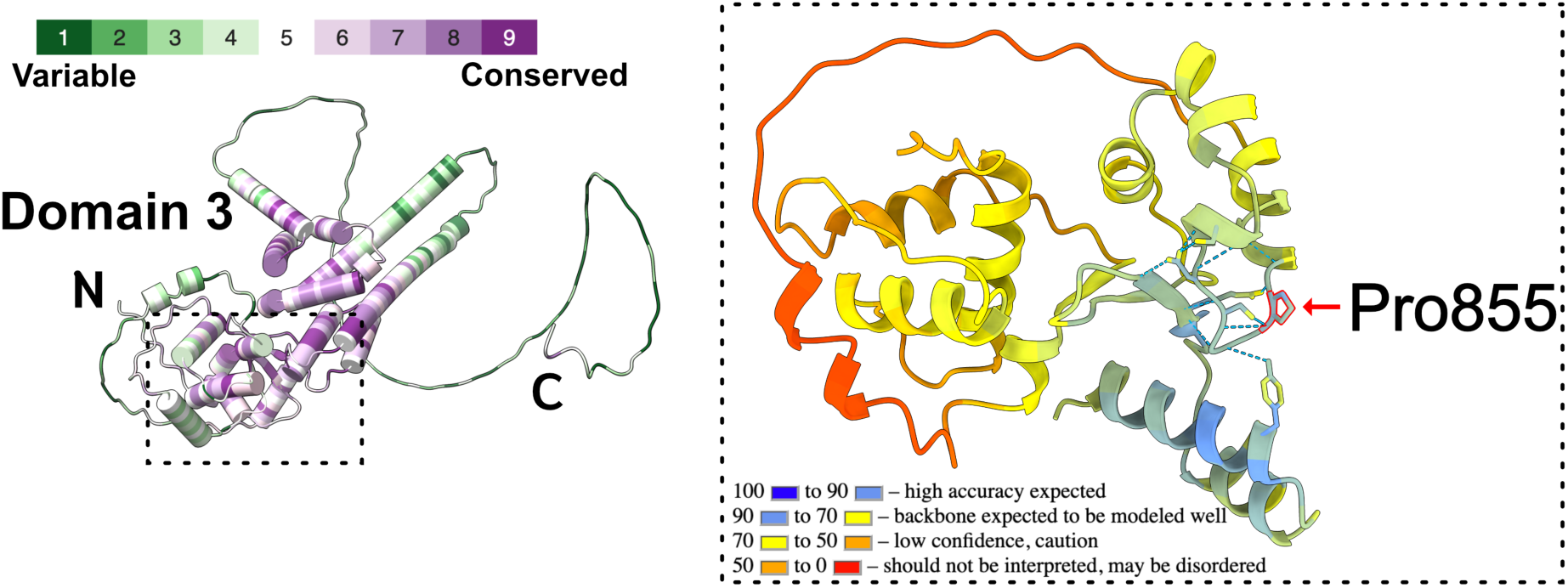
The *nov-1* (Pro855->S) mutation in Tsugeki *et al*., 2009 is in a conserved structural region of an N-terminal domain in NOV. Domain 3 of NOV with ConSurf conservation score coloring based on alignments of NOV homologs in eukaryotes projected on the AF3 predicted structure of AtNOV. The highlighted box shows the area where the *nov-1* Pro855S mutation is localized, colored using ChimeraX pLDDT coloring. Putative hydrogen bonds close to the residue are shown with blue dotted lines.

**Supplementary Figure 5:**
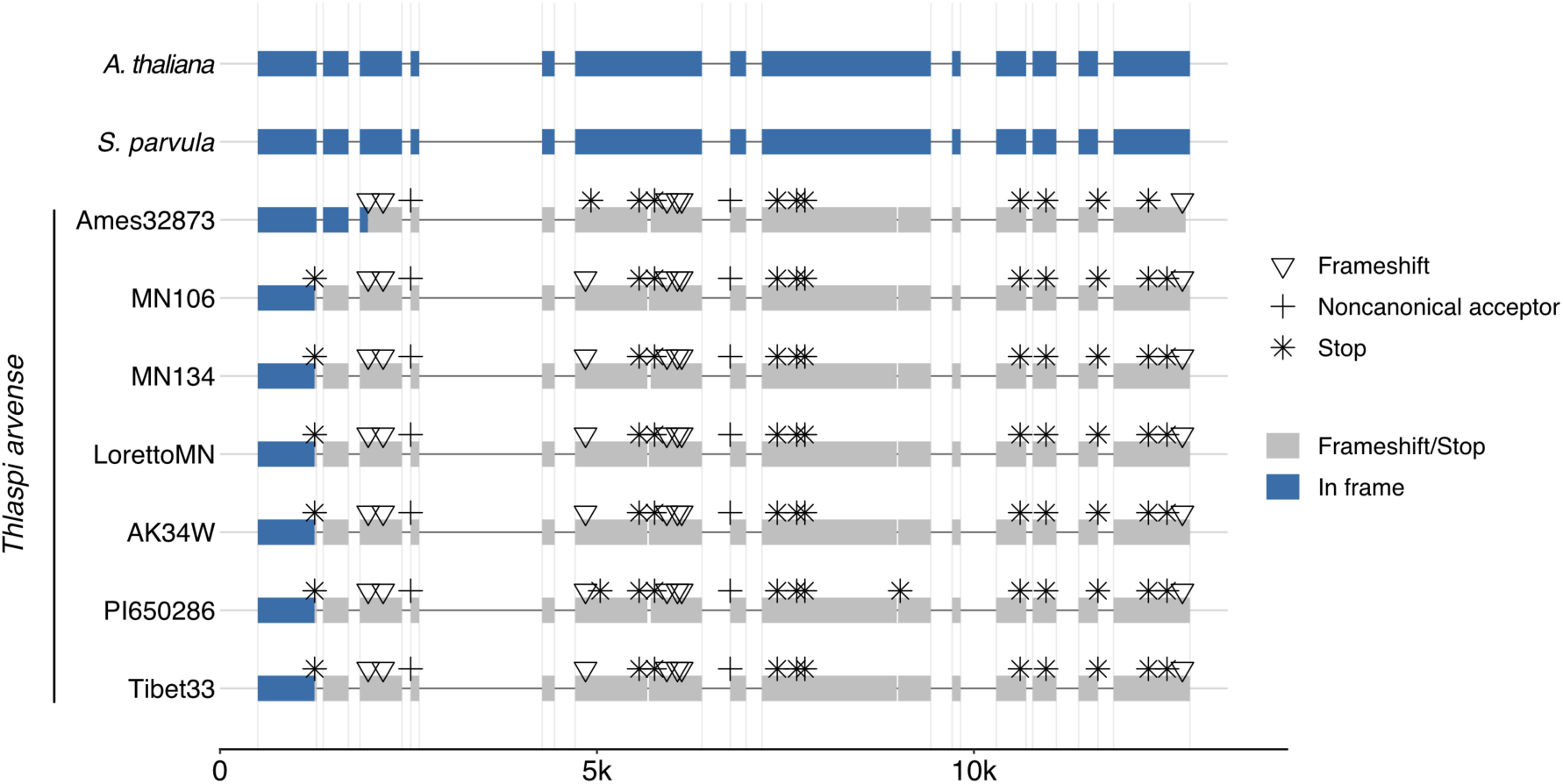
*NOV* is pseudogenized in *Thlaspi arvense*. *NOV* is pseudogenized in all *Thlaspi arvense* accessions analyzed. Regions in frame compared to the *A. thaliana NOV* coding sequence (AT4G13750) are highlighted in blue, while truncated and out of frame regions are highlighted in grey. Frameshifts are marked with triangles, premature stop codons are marked by stars, and noncanonical splicing (acceptor) junctions are marked with plus. *T. arvense* genomes are from the PennyPan project (Bird et al., 2026), and *Schrenkiella parvula* was used as a closely related outgroup to *T. arvense* (Oh et al., 2014).

**Supplementary Figure 6:**
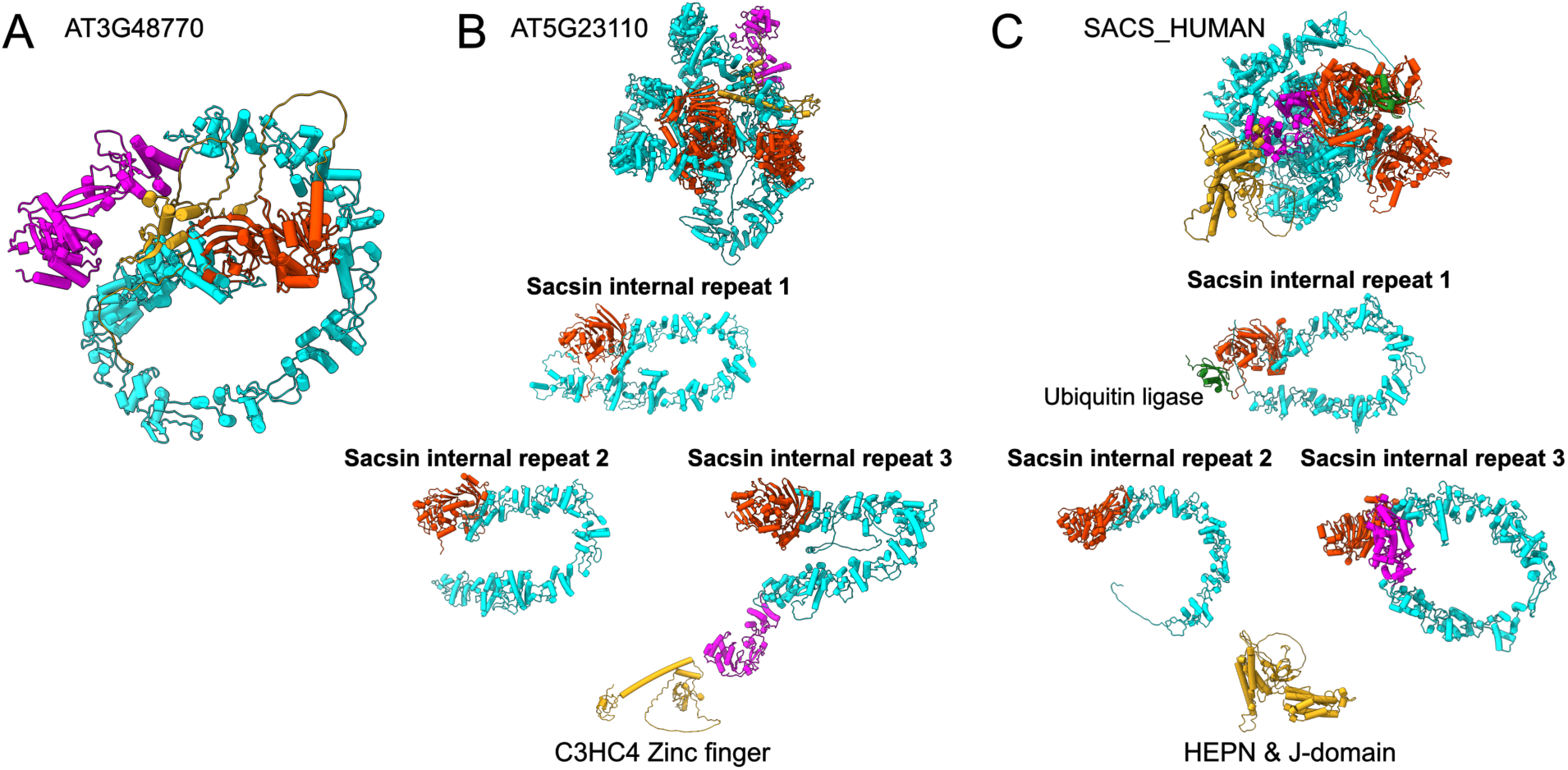
Sacsin-like proteins in *A. thaliana*. **A** Full AlphaFold3 predicted structure of AT3G48770 shows it consisting of a SACS/Tiamat-like ATPase region (orange) and a solenoid repeat region (cyan) as in Tiamat and NOV. This protein has a Zinc finger BED-type domain (PF02892) (gold) at the N-terminus, and the conserved beta-sheet fold (magenta) at the C-terminus found in NOV, Tiamat and other homologs, as well as AT5G23110 and Sacsin. **B** Full AlphaFold3 predicted structure of AT5G23110 shows that it consists of a similar solenoid repeat region as in AT3G23110, with three tandem repeats of the sacsin internal repeat supradomain - which consists of the ATPase domain (orange) and solenoid repeat region (cyan). The third repeat has a beta-sheet fold at the C-terminus, followed by a Zinc-finger C3HC4 domain (PF13920) (gold). Homologs of this protein are often classified as sacsin in automatic annotations, but Ubiquitin ligase, HEPN and J-domains are absent. **C** Full AlphaFold3 structure of Homo sapiens SACS consisting of a Ubiquitin ligase (green) with three Sacsin internal repeat domains (orange and cyan) and a beta-sheet fold (magenta) at the C-terminus of the last repeat, followed by the HEPN and J-domains (gold).

## Notes

### Competing Interest Statement

The authors have declared no competing interest.

